# Fractional diffusion theory of balanced heterogeneous neural networks

**DOI:** 10.1101/2020.09.15.297614

**Authors:** Asem Wardak, Pulin Gong

## Abstract

Interactions of large numbers of spiking neurons give rise to complex neural dynamics with fluctuations occurring at multiple scales. Understanding the dynamical mechanisms underlying such complex neural dynamics is a long-standing topic of interest in neuroscience, statistical physics and nonlinear dynamics. Conventionally, fluctuating neural dynamics are formulated as balanced, uncorrelated excitatory and inhibitory inputs with Gaussian properties. However, heterogeneous, non-Gaussian properties have been widely observed in both neural connections and neural dynamics. Here, based on balanced neural networks with heterogeneous, non-Gaussian features, our analysis reveals that in the limit of large network size, synaptic inputs possess power-law fluctuations, leading to a remarkable relation of complex neural dynamics to the fractional diffusion formalisms of non-equilibrium physical systems. By uniquely accounting for the leapovers caused by the fluctuations of spiking activity, we further develop a fractional Fokker-Planck equation with absorbing boundary conditions. This body of formalisms represents a novel fractional diffusion theory of heterogeneous neural networks and results in an exact description of the network activity states. This theory is further implemented in a biologically plausible, balanced neural network and identifies a novel type of network state with rich, nonlinear response properties, providing a unified account of a variety of experimental findings on neural dynamics at the individual neuron and the network levels, including fluctuations of membrane potentials and population firing rates. We illustrate that this novel state endows neural networks with a fundamental computational advantage; that is, the neural response is maximised as a function of structural connectivity. Our theory and its network implementations provide a framework for investigating complex neural dynamics emerging from large networks of spiking neurons and their functional roles in neural processing.

## I. INTRODUCTION

Networks composed of a large number of interacting units are ubiquitous in physical, biological, financial and ecological systems [1–4]. In these systems, interactions of units give rise to complex network dynamics with great fluctuations occurring at multiple spatial and temporal scales [1–4]. Understanding such complex dynamics is a long-standing topic of common interest in these diverse systems. In neuroscience, neural networks of many interacting neurons likewise exhibit great fluctuations in their firing activity [5–7]; such fluctuations provide a critical window into understanding the working mechanism of neural systems. The classical theoretical framework for understanding how complex neural dynamics emerge is based on homogeneous, randomly coupled networks with balanced excitation and inhibition [8–12]. Such balanced network models have successfully reproduced many experimental observations such as the high irregularity of neural firing activity and broad distributions of firing rates, and have been analysed using dynamical mean field models with the assumption of homogeneous Gaussian noise as synaptic inputs, often referred to as the normal diffusion theory [8–10]. The standard Fokker-Planck formalism is then used in this theoretical framework to obtain the key statistical properties of complex neural dynamics. Virtually all existing balanced neural network models, including the classical ones [8–10] and their recent extensions incorporating heterogeneous degrees [13–15], distance-dependent connectivity [16–18], and alternative neuron models such as conductance-based neurons [19, 20], have been studied by applying such normal diffusion-based formalisms.

Recent experimental studies, however, have been increasingly demonstrating that heterogeneous, non-Gaussian properties are ubiquitous for both neural connectivity and neural activity dynamics. For instance, it has been found that synaptic connection strengths [21, 22] and the number of synaptic connections [23] exhibit heavy-tailed distributions with non-Gaussian features. Furthermore, it has been empirically found that neural firing activity has super-Poisson dynamics [7], with great heterogeneity and fluctuations occurring at multiple scales [24–26], and that membrane potentials fluctuate far from the firing threshold [27]. Some of these properties are not necessarily incompatible with Gaussian input statistics: non-Gaussian, lognormal distributions of synaptic strengths lead to the classical, normal diffusion theory in the large network limit. Suitable assumptions on the classical theory such as adaptation can be incorporated into individual modeling studies, in order to reproduce individual phenomena such as large fluctuations of membrane potentials [28]. Nevertheless, the fundamental problems of whether and how the non-Gaussian features of neural dynamics can be explained in a unified theoretical framework, and the mechanistic relationships between these neural dynamics and heterogeneous connectivity, remain unclear.

Here, extending the classical theory of balanced networks, we develop a novel theory explaining how the non-Gaussian features of complex neural dynamics emerge from biologically realistic, heterogeneous neural networks. Based on balanced spiking neural networks with heterogeneous connection strengths, our analysis reveals that synaptic inputs in such heterogeneous networks possess heavy-tailed, Lévy fluctuations, a type of fluctuation typical for non-equilibrium systems with many interacting units [29–31]. This thus leads to a relation between complex neural dynamics and fractional diffusion formalisms developed for studying non-equilibrium physical systems [32–37]; correspondingly, we refer to this as the fractional diffusion theory of balanced neural networks. In our fractional theory, we mathematically demonstrate how these biologically realistic neural networks lead to membrane potentials undergoing fractional, Lévy diffusion instead of the normal, Brownian diffusion arising in homogeneous networks. By uniquely accounting for the Lévy leapovers caused by the great fluctuations of spiking activity, we develop an analytically tractable fractional Fokker-Planck equation with absorbing boundary conditions. The fractional theory leads to an exact description of the activity states of balanced heterogeneous neural networks; we verify our theory in spiking neural network models. We also demonstrate how heterogeneous firing rate distributions can be analytically incorporated into our fractional diffusion theory.

Based on this fractional diffusion theory of heterogeneous neural networks, we identify a new network state, which is fundamentally different from the asynchronous irregular state and the synchronous irregular state in the conventional models [9, 10, 38]. Due to the fractional nature of this new state, we term it the fractional state. This state exhibits rich nonlinear response properties and can explain a variety of complex neural dynamics at the individual-neuron and circuit levels, which otherwise would be treated in isolation in previous studies. In this state, membrane potential resides far from the threshold, exhibiting heavy-tailed, skewed distributions, quantitatively consistent with neurophysiological recordings in the visual cortex of awake monkeys [27]. Neural firing activity exhibits great variability with firing rate fluctuations, as observed in [7, 24, 26]. The collective firing rate fluctuations of neural networks happen at multiple scales in a scale-free manner, as widely observed in experimental studies [25, 39]. We further demonstrate that in this fractional state, the firing rate for a given external input is maximized; that is, the neural response is maximised as a function of structural connectivity. Our fractional diffusion framework not only provides a novel, unified account of a variety of the key features of neural dynamics, but can also be applied to understand how complex dynamics emerge from other non-equilibrium systems with large numbers of interacting units.

## II. HETEROGENEOUS NETWORK MODEL

We consider a generalization of the classical homogeneous networks which have been extensively studied over the past two decades [8–11, 38, 40]. Our circuit model consists of *N*_*E*_ excitatory and *N*_*I*_ inhibitory neurons. Each neuron receives *C*_*E*_ and *C*_*I*_ connections from excitatory and inhibitory neurons, respectively, and *C*_ext_ from excitatory neurons outside the network. The membrane potential *V*_*i*_(*t*) of neuron *i* ∈ {1, *…, N*_*E*_ + *N*_*I*_} satisfies the equation for a leaky integrate-and-fire neuron [10],

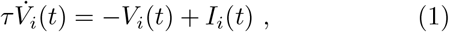

where *τ* is the integration time, and *I*_*i*_(*t*) denotes the total synaptic current to neuron *i*. The current is modeled as a weighted sum of delta functions representing spikes from incoming neurons,

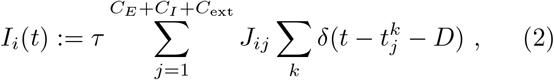

where *J*_*ij*_ is the synaptic connection strength from neuron *j* to neuron *i*; 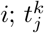 is the emission time of the *k*th spike of neuron *j*; and *D* is the transmission delay. The spikes of external neurons are modeled as independent excitatory neurons with Poisson spike trains at rate *v*_ext_. The *i*th neuron spikes when *V*_*i*_(*t*) reaches the firing threshold *θ*, resetting the membrane potential to the reset potential *V*_*r*_ after a refractory period *τ*_rp_ during which the membrane potential is unresponsive to input.

In previous studies, heterogeneous heavy-tailed distributions of synaptic weights have been fitted to lognormal distributions [21–23]. Because of the central limit theorem, in the thermodynamic limit of large network size, the dynamics of neural networks with lognormal coupling weights should be equivalent to those under the homogeneous Gaussian hypothesis (Fig. 1(a)), yielding the classical asynchronous state with Poisson-like spikes. To theoretically explore the effect of heterogeneous connectivity on network dynamics, we assume that the heavy-tailed distributions of synaptic weights are power laws, the same assumption made in [41] to theoretically study the effect of heterogeneous connectivity on neural dynamics near the edge of chaos. Recent experimental studies have found the existence of power-law synaptic strengths in drosophila whole brain data [42], but note that these are not mammalian brain data. In our heterogeneous network model (Fig. 1(b)), heavy-tailed distributions of outgoing synaptic weights *J*_*ij*_ are approximated by power law distributions. This power-law assumption, as demonstrated in the following sections, leads to a novel fractional diffusion framework that accounts for a variety of non-Gaussian properties of neural dynamics such as the infrequent large fluctuations of membrane potential and the power-law scaling of the Fano factor of spike counts. The relationship between the outgoing, statistically independent synaptic strengths in the excitatory (*J*_*E*_) and inhibitory (*J*_*I*_) populations is ⟨*J*_*I*_⟩ = *g* ⟨*J*_*E*_⟩ = *gJ*, where *g* is a balance factor describing the relative strength of inhibitory inputs, and *J* := ⟨*J*_*E*_⟩ = ⟨*J*_ext_⟩.

**FIG. 1.**
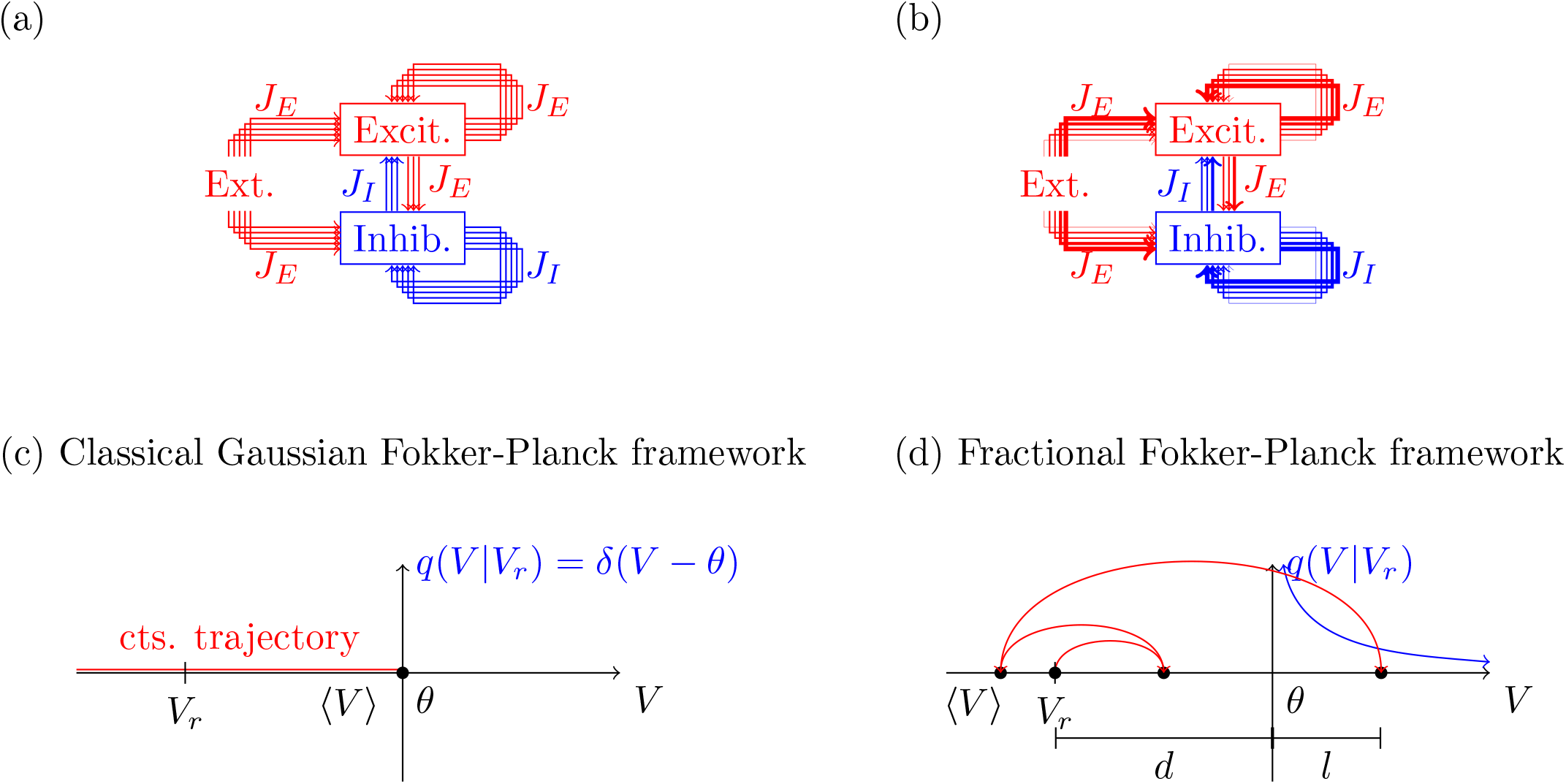
Schematic of the stochastic formalisms of homogeneous and heterogeneous networks. (a) The classical formalism begins with a network model where the connection strengths *J*_*ij*_ are constant (or drawn from a Gaussian distribution), denoted by the lines of *J*_*E*_ and *J*_*I*_ with constant thickness. The classical theory leads to a Fokker-Planck framework (c), where the membrane potential *V* resides close to the threshold *θ* with Gaussian fluctuations; it thus never crosses the threshold without passing through it (first arrival equaling first passage) and so the sink is a delta function at the threshold. (b) The network is strongly heterogeneous and the connection strengths are drawn from a power-law distribution, denoted by the lines with different thicknesses, leading to a fractional Fokker-Planck formalism (d) whereby the membrane potential resides far from the threshold and has occasional large fluctuations, due to the presence of large jumps in the trajectory. This leads to leapovers *l* over the boundary which must be accounted for correctly in the fractional Fokker-Planck equation by using the first passage leapover density as a spatially extended sink.

### A. Fractional diffusion formalism for heterogeneous networks

As in the classical normal diffusion theory for homogeneous networks [8–11, 13–20], we consider the dynamics of our heterogeneous network in the limit of large network size. By using the generalized central limit theorem [43], we rigorously prove that the total synaptic input *I*(*t*) to an individual neuron in the network has non-equilibrium Lévy fluctuations, in the limit of large network size, if and only if the distributions *J*_*r*_ of synaptic strengths outgoing from population *r* ∈ {*E, I*, ext} have a power-law tail (see detailed derivation in Appendix A). For all other distributions, however, the classical Gaussian fluctuations are recovered.

The total synaptic current *I*(*t*) to a given network neuron is thus expressed as the sum of an average term with a fluctuating Lévy noise term,

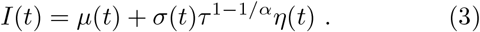

The noise term *η*(*t*) is driven by an *α*-stable Lévy process *L* = *L*(*α, β, γ*_*L*_, *D*_*L*_) [44] with Lévy stable index 1 *< α* ≤ 2, skew parameter *β* given by Eq. (A23),

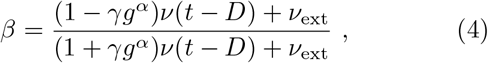

center *γ*_*L*_ = 0, and scale *D*_*L*_ determined by the tails of the connection strength distributions *J*_*r*_. The classical diffusive case is recovered when *α* = 2, at which point the Lévy process *L*(2, *β*, 0, *D*_*L*_) = 𝒩(0, 2*D*_*L*_) reduces to a Gaussian process. The term *v*(*t*) represents the mean population firing rate at time *t*. The mean input term *µ*(*t*) is a sum of internal and external inputs to the neuron, while *σ*(*t*) is the fluctuation in the sum of the internal and external inputs (see Appendix A),

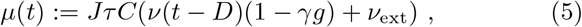

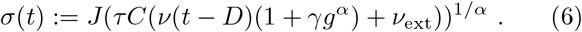

Here, for clarity, we have taken *N*_*I*_ = *γN*_*E*_, *C* := *C*_*E*_ = *C*_ext_, and *C*_*I*_ = *γC*, where *γ* = 0.25 is the ratio of inhibitory to excitatory neurons; the general equation without these constraints is given by Eq. (A21). As in previous studies on the classical normal diffusion theory [13, 14], this formalism can be extended to compute population firing rate distributions *v*_*r*_(*t*); a mean-field method for this under the fractional formalism is detailed in Appendix B.

Using the relation *dL* = *η*(*t*)*dt*, the dynamics of the neurons in Eq. (3) can be expressed using a Langevin-style stochastic differential equation,

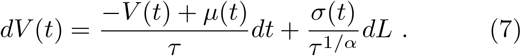

The membrane potential can thus be regarded as a particle driven by Lévy noise, moving in a quadratic external well centered at *µ*(*t*).

The external input *v*_ext_ is expressed in units of *v*_thr_ = *θ/*(*CJτ*), which is the external frequency required for the mean input *µ*(*t*) to reach threshold in the absence of recurrent feedback. When the populations are balanced (*g* = *γ*^−1^ = 4), we obtain the balance condition in the sense of [15],

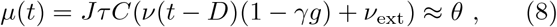

which is satisfied for (*g, v* _ext_) = (*γ*^−1^, *v* _thr_). In the following we focus on balanced heterogeneous networks, although the expressions are applicable more generally as *g* and *v*_ext_ vary.

## III. FRACTIONAL FOKKER-PLANCK EQUATION AND CONSTRUCTION OF BOUNDARY CONDITIONS

By applying the theory of Lévy processes, we obtain the membrane potential probability density *P* (*V, t*) corresponding to Eq. (7), which satisfies a fractional Fokker-Planck equation [44]

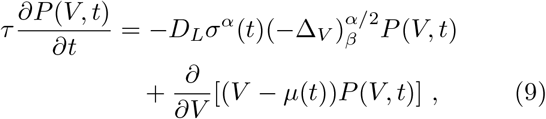

where

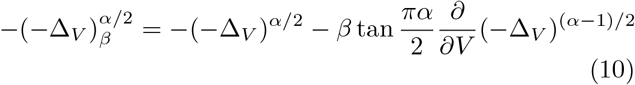

is the skewed fractional Laplacian, and

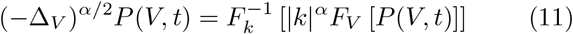

is the fractional Laplacian, defined as a Fourier multiplier. The right hand side of Eq. (9) consists of a fractional diffusive term arising from the fluctuations of the input, along with a drift term from the average part of the input. Since 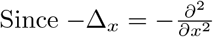, the equation can be rewritten as a continuity equation,

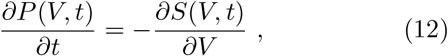

where

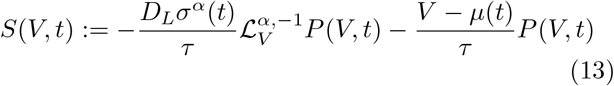

is the probability current through *V* at time *t*, and

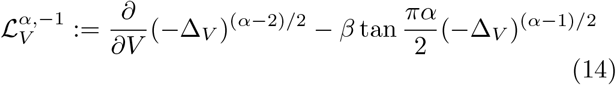

has the property 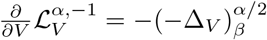.

In order to map the spiking behavior of neurons onto the fractional Fokker-Planck equation (9), the boundary conditions need to be specified at the reset potential *V*_*r*_ and threshold *θ*, along with a normalization condition. In the conventional Fokker-Planck framework, neural spiking has been defined as first arrival to the threshold *θ*. The firing of the neuron thus imposes a point sink term *v*(*t*)*δ*(*V* − *θ*) [19] at the threshold *θ*, where probability density is actively removed with rate *v*(*t*); this is illustrated in the schematic in Fig. 1(c). Neurons exiting the refractory period are injected back into the probability density using a point source term *v*(*t* − *τ*_rp_)*δ*(*V* − *V*_*r*_) at the reset potential *V*_*r*_. This injection happens at a rate equal to the firing rate of the network when those neurons had just fired, *v*(*t* − *τ*_rp_).

In our fractional case, however, the situation is more nuanced due to the discontinuous large jumps of the membrane potential *V* (*t*) inherent to Lévy processes, which are absent in the continuous Gaussian trajectories. This allows the membrane potential to jump over the boundary without arriving at it; as a result, the first passage of the membrane potential *V* (*t*) through the threshold *θ* is no longer necessarily equivalent with the first arrival of *V* (*t*) at *θ* (see Fig. 1(d)) [35]. This poses a problem with using the conventional Fokker-Planck derivation of absorbing boundary conditions in the fractional case. If neural spiking is still defined as first arrival to the threshold *θ*, then the membrane potential *V* (*t*) may spend time above the threshold before firing, which is unphysical in our network model. In order for the neuron to correctly fire whenever the membrane potential is found to be above threshold, one must define neural spiking to be first passage rather than first arrival with respect to the threshold *θ*. Since first arrival equals first passage in the conventional Gaussian case, the definition of neural spiking as first passage is physically consistent in both the conventional and fractional cases.

When neural spiking is defined as first passage, the sink term becomes spatially extended in the fractional case, equal at each point *V* ≥ *θ* to the probability that the membrane potential *V* (*t*) lands at *V* at the time of first passage through the threshold *θ*. In the absence of resetting, the sink term is determined by the first passage leapover density [45]. The Langevin equation (7) for the membrane potential in the balanced condition (*g, v*_ext_) = (*γ*^−1^, *v*_thr_) corresponds to Lévy motion in a quadratic external well with absorbing boundary at its extremum. In this case, the first passage leapover density *q*(*V, t*) can be obtained from a variable transformation of the leapover in the free case [45, 46],

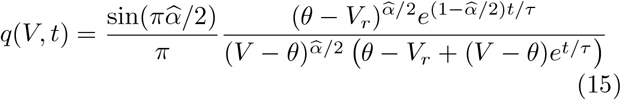

for *V* ≥ *θ* where

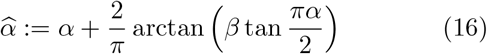

is related to the positivity parameter *ρ* [47–49] by the identity 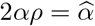.

When neurons exiting the refractory period are also considered, the canonical absorbing sink [46] becomes *v*(*t*) *_*t*_ (*p*_FP_(*t*)*q*(*V, t*)), where *_*t*_ denotes temporal convolution, and *p*_FP_(*t*) is the first passage time density of a particle driven by Lévy motion in a quadratic external well out of (−∞, *θ*) starting at *V*_*r*_. As time passes, an expanding region is established around the reset potential *V*_*r*_, characterized by the nonequilibrium stationary state [50]. Within this region, the first passage time density in the convolution may be regarded as a delta function *p*_FP_(*t*) = *δ*(*t* − 1*/v*(*t*)) [46]. After a large amount of time has passed, the sink term everywhere approaches the expression *v*(*t*)*q*(*V*, 1*/v*(*t*)). The analytical results in this work are expressed in terms of *q*(*V, t*) in order to account for the unbalanced case (*g, v*_ext_) ≡ (*γ*^−1^, *v*_thr_) when its corresponding first passage leapover density *q*(*V, t*) becomes known in the statistical physics literature [32–37, 45–50]. Since neurons are always reset to the same reset potential *V*_*r*_, the point source term *v*(*t* − *τ*_rp_)*δ*(*V* − *V*_*r*_) due to neurons exiting the refractory period remains unchanged as in the classical Gaussian case.

Next, by the definition of the integrate-and-fire neuron, *P* (*V, t*) = 0 for *V > θ*, and thus by continuity

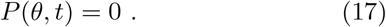

Since *P* (*V, t*) is a probability distribution, it is integrable (so its spatial Fourier transform 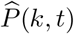 is smooth):

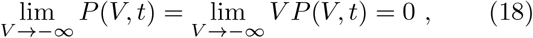

and satisfies the normalization condition

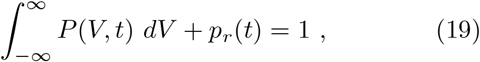

where 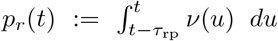 is the probability that the neuron is refractory at time *t*. Incorporating these boundary conditions into Eq. (9) yields a novel, analytically tractable fractional Fokker-Planck equation for characterizing motion in the presence of absorbing barriers,

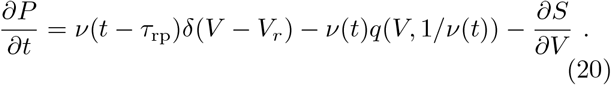

This equation is novel in that previous approaches to incorporating absorbing barriers in the fractional Fokker-Planck equation required the truncation of the integral form of the fractional Laplacian, preventing standard analytical methods from being used [35]. Here, however, the absorbing boundary is fully encapsulated in the extended sink term, allowing the fractional Laplacian to remain untouched and hence permitting the usage of standard techniques for its analytical solution.

## IV. NETWORK DYNAMICS REVEALED BY THE FRACTIONAL FOKKER-PLANCK FORMALISM

We now show that Eq. (20) can be solved under the condition *∂P/∂t* = 0, in order to show the stationary network dynamics of the membrane potential distribution *P*_0_(*V*) and firing rate *v*_0_, and uncover the existence of a novel state at which the neural response is structurally optimal. This can be done by moving into the Fourier domain and solving the resultant ordinary differential equation, as detailed in Appendix C. We denote the variables corresponding to the time-independent solution of Eq. (20) by the addition of a 0 to their subscripts, and the omission of their time arguments; the stationary membrane potential distribution *P*_0_(*V*) given by Eq. (C12) is

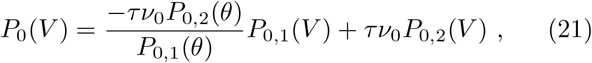

where *P*_0,1_(*V*), *P*_0,2_(*V*) are defined in Eqs. (C13, C14). Although the fractional diffusion theory can also be derived using other neuron models, for which the first term on the right hand side of Eq. (1) becomes an arbitrary function of *V*_*i*_, we have chosen the theoretically simplest neuron model in order to focus on the heavy-tailed heterogeneity. The stationary firing rate *v*_0_ can be obtained by solving the implicit equation (C19), which is

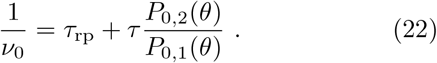

This generalizes the classical result for the mean first passage time of an IF neuron with random Gaussian inputs to the Lévy case [10, 51]. We then directly calculate the stationary membrane potential distributions (Eq. (21)), and numerically solve the stationary firing rate equation (22) using Mathematica [52] and the biologically realistic parameter sets detailed in Appendix D.

Figure 2(a) shows the stationary firing rate *v*_0_ (see Eq. (22)) as a function of the Lévy stable index *α*, all other parameters such as the external firing rate being kept constant. As *α* decreases from the classical Gaussian value *α* = 2, the probability for the membrane potential *V* to encounter a sequence of large jumps towards the threshold *θ* increases. This causes an increase to the firing rate, despite the external input rate remaining constant. However, for low *α* close to *α* = 1, the probability for a large jump of the membrane potential away from the threshold, from which it would be difficult to recover with even moderately sized jumps, begins to become significant, causing a reduction in the firing rate. These two counteracting forces on the firing rate, arising from the strong heterogeneity of synaptic strengths, are balanced around the value *α* = 1.2, representing the maximum response in activity for a given external input rate; this state is referred to as the fractional state. To highlight the differences caused solely by a change in structural connectivity, we have obtained all values in Fig. 2 using the biologically realistic parameter set (see Appendix D). Here, the firing rate in the classical Gaussian regime (*α* = 2) is around 18 Hz, while in the fractional regime (*α* = 1.2), the firing rate rises to 54 Hz, a three-fold increase due solely to the heterogeneity of the network. These results thus reveal that heterogeneous neural coupling can have profound functional advantages in neural networks.

**FIG. 2.**
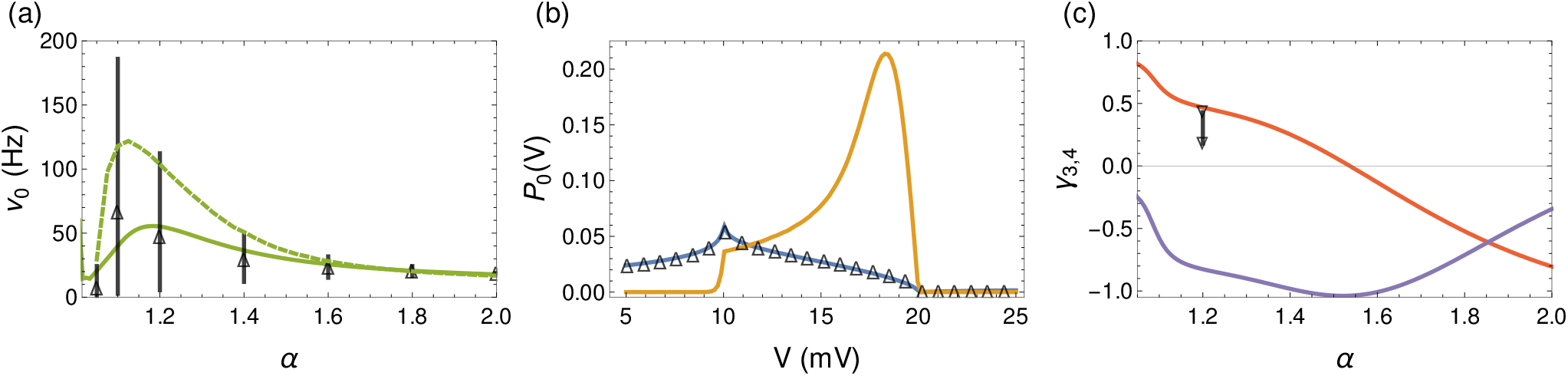
Stationary states vary with the fractional Lévy index *α*. (a) The stationary firing rate *v*_0_ as a function of *α* under the homogeneous-rate (green) and mean-heterogeneous (green-dashed) assumptions is maximized around the optimal Lévy range *α* ∈ (1.1, 1.2), which is significantly higher than the firing rate arising in the classical asynchronous state (*α* = 2). The vertical lines represent 5th-95th percentile ranges in the corresponding network firing rate distribution, and the triangles show their means. (b) The membrane potential distributions *P*_0_(*V*) for the fractional state *α* = 1.2 (blue) and the Gaussian state *α* = 2 (yellow) under the homogeneous-rate assumption agrees with the network values (triangles). (c) The skewness *γ*_3_ (red) and excess kurtosis *γ*_4_ (purple) of the membrane potential distribution under the homogeneous-rate assumption as *α* varies; the vertical line shows the decrease to the skewness of the network distribution when adding constant input to the network.

At this point of maximal response, the membrane potential rests far from the threshold (Fig. 2(b), blue line) in contrast to the classical case, where the membrane potential hovers close to the threshold (Fig. 2(b), yellow line). To further quantify this difference and compare with experimental results on membrane potential residuals, we calculate the skewness *γ*_3_ of the membrane potential distribution between the reset potential and threshold, which is around −0.7 and 0.5 in the Gaussian and fractional cases respectively (Fig. 2(c), red line). These properties of membrane potential predicted by our theory are consistent with recent whole-cell *V*_*m*_ measurements from the visual cortex of behaving monkeys [27], in which it has been demonstrated that *V*_*m*_ is far from threshold during spontaneous activity, and its distribution is skewed positively with median value *γ*_3_ = 0.72 [27]. Although leaky integrate-and-fire neurons have a hard threshold, a feature not present in other spiking neuron models such as the exponential integrate-and-fire neuron, they remain effective in capturing the subthreshold properties of membrane potentials.

The zero density of the membrane potential distribution above threshold in Fig. 2(b) demonstrates that the absorbing boundary condition has been set up properly; when the classical first arrival point sink *v*(*t*)*δ*(*V* − *θ*) is instead used for the boundary condition (see the schematics in Figs. 1(c, d)), the membrane potential distribution in the fractional case becomes nonzero above the threshold point *θ* [35], representing the trajectories of the membrane potential which jump over the threshold without arriving at it.

## V. SPIKING NETWORK IMPLEMENTATION

To verify our fractional diffusion theory and further illustrate that it can unify a range of neural dynamics [7, 24, 27], we next numerically investigate heterogeneous balanced neural networks with the distributions of heavy-tailed coupling weights approximated as power-law functions (as detailed in Sec. II).

In line with the findings in [15], we have found that when the synaptic strengths are drawn at random from their distributions (see Eq. (A15) and Appendix D), the per-neuron balance between excitation and inhibition is hindered, leading to the division of the network into silent and saturated neurons. In order to mitigate these effects, without violating the statistical independence of inputs across different populations, we ensure that the overall balance between the sums of excitatory and inhibitory connection strengths to a given neuron is sufficiently close to *g* = *γ*^−1^ = 4, so that

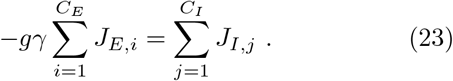

This tight balance condition can be naturally recovered using neural mechanisms such as those discussed in [15]. In our network model, we ensure tight balance for each neuron by drawing the ensembles of excitatory weights once, and then redrawing the ensembles of inhibitory weights, until the absolute ratio between the excitatory ensemble sum and the redrawn inhibitory ensemble sum is to within *ε* = 10^−3^ of 1*/gγ*. Importantly, we recover this balance while maintaining the heterogeneity in input connectivity.

We use the spiking neural network package NEST [53] to simulate a network of *N*_*E*_ excitatory and *N*_*I*_ inhibitory neurons; biologically realistic parameter sets (Appendix D) are used to highlight the effect of fractional, Lévy noise compared with the classical, Gaussian asynchronous irregular regime.

To represent the behavior of the fractional regime, we have chosen the value *α* = 1.2, around which the firing rate response is maximized (see Fig. 2(a)); the essential behavior of the fractional regime with respect to neural variability, however, can be seen for any *α* sufficiently far from the Gaussian value *α* = 2. In particular, numerical simulations of network firing rates at various values of *α* (Fig. 2(a), triangles and vertical lines) are generally consistent with analytical predictions and demonstrate that firing rate distributions become very wide across neurons in the network in the fractional regime, which are lognormal as we explain below (see Fig. 3(c)).

**FIG. 3.**
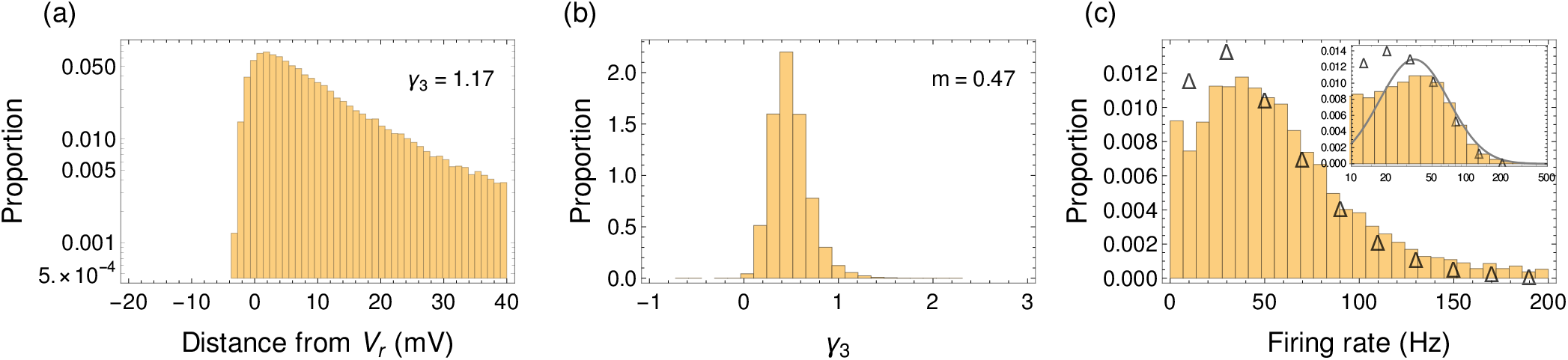
Per-neuron properties of the stationary fractional state (*α* = 1.2). (a) The distribution of distances of the mean membrane potential from the reset potential *V*_*r*_ across neurons has skewness *γ*_3_ = 1.17. (b) The distribution of membrane potential skewness across neurons has median value *γ*_3_ = 0.47. (c) Firing rate distribution using the Monte-Carlo method applied to the mean-field formalism in Appendix B (histogram), and corresponding network distribution (triangles); lognormal fit in inset.

To validate the prediction of our fractional diffusion theory, we compute the membrane potential distribution across all neurons in the network (Fig. 2(b), triangles), which is in excellent agreement with the theoretical stationary state values. To directly compare our results with experimental observations, we compute the per-neuron properties of membrane potentials in the fractional regime, as shown in Figs. 3(a,b). The heterogeneity in connectivity leads to a wide distribution of means and skewness of membrane potentials across neurons in the network; these heterogeneous properties across neurons are consistent with those measured in [27] (see their Figs. 2(e,f)). Since classical, randomly connected networks with heterogeneous in-degrees are expected to have a distribution of skewness, it would be interesting to study how this skewness can be compared with quantitative data as has been done in this study.

To demonstrate how the activity of the network may be switched between the fractional state and one with more Gaussian features, as observed experimentally [27], we perform network simulations (*α* = 1.2) by adding constant external input of larger magnitude. The peak of the network membrane potential distribution flattens, with its skewness decreasing from around 0.5 to 0.1 (Fig. 2(c), black), largely comparable with the decrease in skewness observed in [27] (see their Fig. 3(d)]) after stimulus onset. This change can be analytically explained using the fractional theory in Appendix A, by modeling the external input as a subpopulation with heterogeneity index *α* = 2, while maintaining the *α* = 1.2 index of the internal network subpopulations.

Next, we calculate the distribution of firing rates across neurons in the heterogeneous network; in our fractional regime, the network displays a lognormal firing rate distribution (Fig. 3(c), triangles), negatively skewed as in [12, 54], but with the in-degrees *C*_*E*_, *C*_*I*_, *C*_ext_ remaining constant throughout the network. The novel mechanism behind this heterogeneous, lognormal firing rate behavior is the highly variable Lévy input that each neuron receives, arising from the independent, heterogeneous structure of the synaptic strengths; those neurons with a weak ensemble of synaptic strengths fire least often, while neurons with strong synaptic strength ensembles have a highly tense excitation-inhibition balance, and consequently have the highest firing rates. The skewed property of the distribution arises from the existence of a membrane time constant in the network, as noted by previous studies [12]. Classical models can robustly generate lognormal firing rate distributions by using randomly connected networks, such as Erdős-Rényi networks with constant connection probability [12–14], without requiring a tight balance condition such as Eq. (23). This is because the two sides of Eq. (23) differ by at most a Gaussian quantity in these classical models, instead of one with a power-law tail in the heterogeneous case without tight balance. Here, however, we have achieved lognormal firing rates in a model where the in-degree is constant. This is in order to isolate power-law inputs arising from network heterogeneity as a mechanism for lognormal firing rates, in conjunction with a variety of experimental findings described in this section.

To assess the ability of the fractional diffusion theory to reproduce firing rate distributions for strongly heterogeneous networks, we perform Monte-Carlo simulations based on the mean-field formulation of heterogeneous firing rates under the fractional theory, in Appendix B, which proposes an additional quenched source of variability *µ*_*q*_ to the mean input *µ*(*t*) (Eq. (B10)):

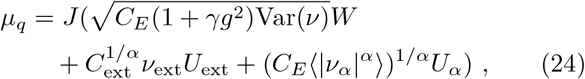

and find that the skewed lognormal behavior of the network is reproduced in essence (Fig. 3(c), histogram). However, as discussed at the end of Appendix B, a significant contribution arises due to local effects (via individual properties of nearby-connected neurons) in the strongly heterogeneous case, which are by assumption dropped in the mean-field formalism. This contribution is clearer when computing fixed points by iterating the Monte-Carlo simulations (Fig. 2(a), green-dashed): the additional correlations discussed in Appendix B serve to restrain the firing rate, but do not change its relative behavior as a function of structural connectivity *α*. In particular, our fractional diffusion formalism is able to capture the network peak of the firing rate at *α* = 1.1 compared to the *α* = 1.2 prediction under the balance condition (23) in the sense of [15].

The spikes of neurons exhibit great variability, as shown in the raster plot (Fig. 4(a)). To quantify such variability, we calculate the coefficients of variation (CV) of the interspike interval (ISI), and find that the distribution of CV is wide in the fractional regime (Fig. 5), ranging between 0.5 and 2.5 with a mean of 1.4. On the other hand, the CVs of the classical homogeneous network with constant in-degree *C*_*E*_ are concentrated around a single CV value (1.0). This variability in CV arises because the ensemble sums of heterogeneous synaptic strengths in the fractional regime, while balanced across different populations, differ on a per-neuron basis, leading to vast fluctuations in firing activity across different neurons. The distribution of CVs in classical networks such as [12] is also broad due to the in-degree heterogeneity.

**FIG. 4.**
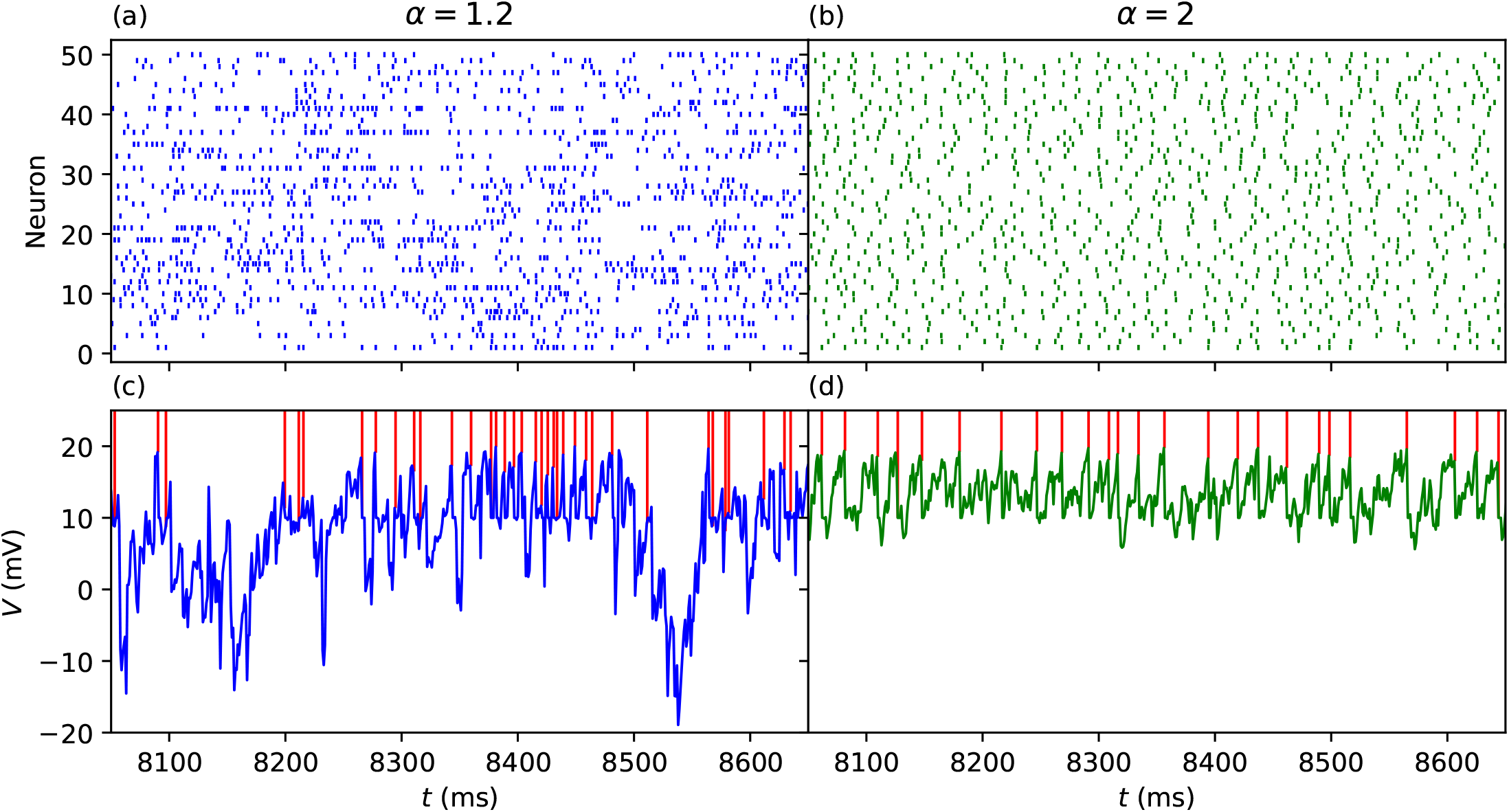
Network activity in the novel fractional (*α* = 1.2) and classical asynchronous (*α* = 2) regimes. Raster plot (top) of 50 randomly chosen excitatory neurons from the network model simulation. Membrane potential time series *V* (*t*) (bottom) of a single network neuron as a function of time *t*.

**FIG. 5.**
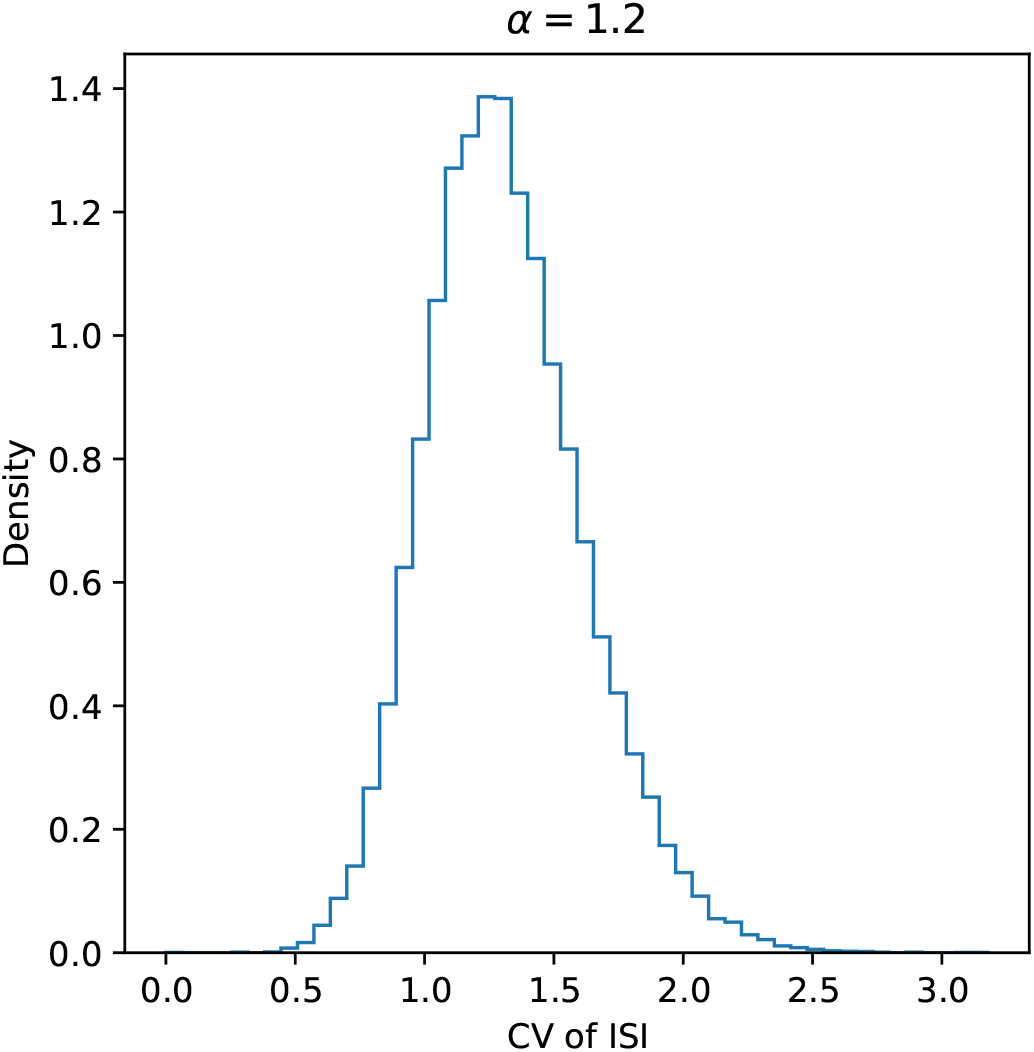
Density histogram of the coefficient of variation (CV) of the interspike interval (ISI) across neurons in the heterogeneous network. The novel fractional regime (blue) has a wide distribution of CVs compared to the classical asynchronous regime.

As shown in Fig. 4(a), firing rates exhibit large fluctuations over long timescales in the fractional regime. To quantify such firing rate fluctuations, as in [24], we compute the Fano factor *F* (*T*) = Var(*N* (*T*))*/* ⟨*N* (*T*) ⟩ with different time window sizes *T*, where *N* (*T*) is the individual neural spike count in time window *T* (Fig. 6). The fractional regime shows that for *T >* 10 ms, the Fano factor increases in a fractal manner as *T* increases, indicating the presence of great firing rate fluctuations. Such fluctuations can actually be fitted as power-law functions as found in [24, 26]. The power-law index of the Fano factor around the optimal parameter *α* = 1.2 is approximately 0.5 (Fig. 6, black dashed line), consistent with experimental findings on the firing activity of cortical neurons [26]. As noted in previous studies [55], the Fano factors in the classical asynchronous state *α* = 2 have the Poisson-like value of unity only when there is no refractory period, with sub-Poisson behavior in the presence of refractory effects; however, the fractional state is able to exhibit a power-law increase in the Fano factor regardless of the existence of refractory effects. This suggests the existence of a phase transition between the Gaussian and fractional regimes, at which the long-term Fano factor switches from zero to infinity respectively [56].

**FIG. 6.**
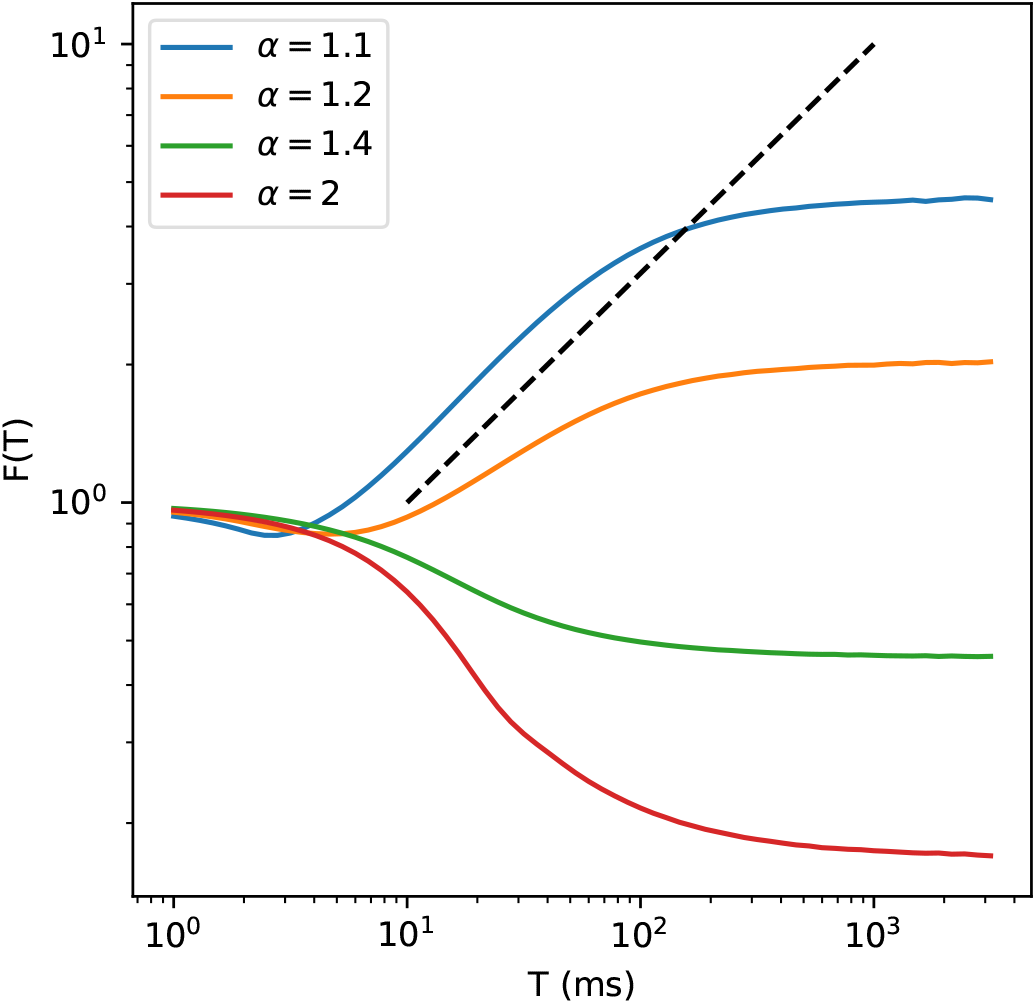
Fano factor *F* (*T*) of single-neuron spiking activity in the heterogeneous network as a function of time window *T*. Black dashed power-law line with exponent 0.5 shown for comparison.

To directly compare with the fluctuations of firing rates as found in [7], we then calculate the variance and mean of the spike count over 1 second windows, with each neuron constituting a single data point. As shown in Fig. 7, the superlinear increase of the spike count variance as a function of its mean is consistent with experimentally observed spike count variability (see Fig. 2 in [7]) in macaque cortical areas V1, V2 and MT. In [7], such fluctuations were modeled based on a phenomenological doubly stochastic model, in which spikes are generated by a Poisson process whose rate is modulated by a slowly fluctuating gain with Gaussian dynamics. However, in our theoretical framework, such firing activity is an emergent, intrinsic property of our biologically realistic, balanced heterogeneous networks. Moreover, the doubly stochastic scenario of [7] leads to a linear scaling of the Fano factor with time window size: from Eq. (3) of [7], 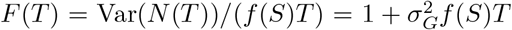, where *f* (*S*) is a function of the stimulus and *G* is a stimulus-independent gain. This is in contrast to the power-law scaling of the Fano factor predicted by our model (Fig. 6).

**FIG. 7.**
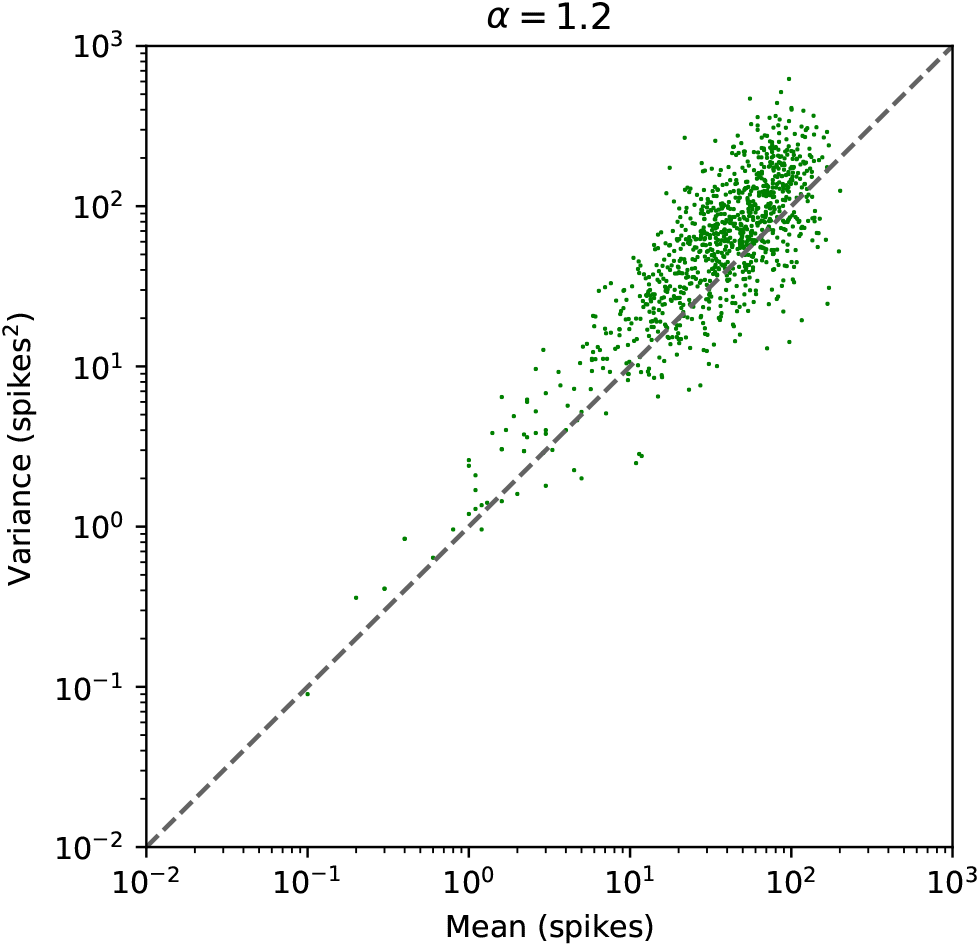
Variance-to-mean relation of the neural responses to a random sample of 1000 excitatory neurons in the balanced heterogeneous network.

We next investigate the fluctuations of population firing rate, which is the ratio between the number of spikes across the network and the time window (1 ms) over which the spikes are counted. We use the analytic Morse wavelet to compute the continuous wavelet transform and obtain the wavelet spectrogram of the population firing rate. In the fractional regime *α* = 1.2, the classical observed frequency band of oscillation deteriorates, with intermittent bursting activity at all frequency scales (Fig. 8(a)). In the classical homogeneous regime *α* = 2, oscillations exist at the characteristic frequency, albeit partially suppressed in the asynchronous irregular state as shown in Fig. 8(a). Frequencies sufficiently far from the characteristic oscillation frequency are fully suppressed compared to the heterogeneous network regime. The power spectrum of the network activity displays 1*/f* - like behavior in the fractional regime (Fig. 8(b)), with a power-law exponent around − 2.0 as observed experimentally [25, 39]. Such 1*/f* -like behavior can also be found using other measures for the activity of the network, such as the local field potential (LFP) calculated as the sum of the total excitatory current and the magnitude of the total inhibitory current [57]. We compute the LFP for the network model with constant external input, which exhibits the 1*/f* property (Fig. 8(b), inset). Such fluctuations of LFPs are thus an intrinsic property of the heterogeneous network; in the conventional balanced network, an external source of fluctuations, such as an Ornstein-Uhlenbeck process with a slowly varying time constant, needs to be incorporated in order to model LFP fluctuations [58]. This scalefree behavior of the firing rate oscillations demonstrates that the classical linear stability analysis is inadequate in the description of neural behaviour around the stationary state for the fractional regime: a strongly nonlinear oscillatory response is observed in the activity with no characteristic oscillation frequency scale. This distinguishes it from both the asynchronous and synchronous irregular classical Gaussian regimes in the sense of [9, 10], where (possibly partially suppressed) oscillations at characteristic frequencies continue to exist.

**FIG. 8.**
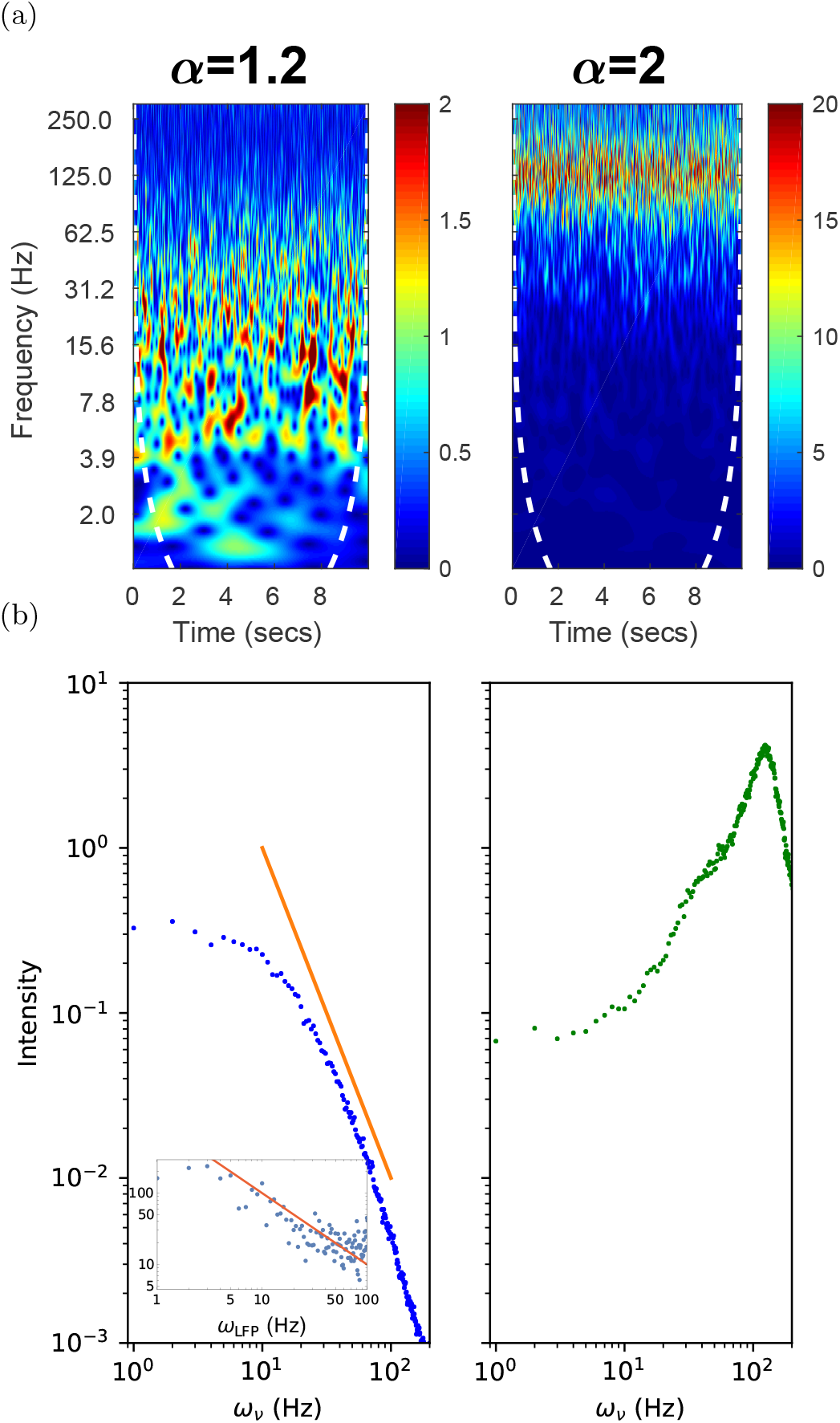
(a) Wavelet spectrogram of network population firing rate as a function of time *t* and oscillation frequency *ω*, for the fractional state, *α* = 1.2 (left), and the classical asynchronous state, *α* = 2 (right). (b) Power spectrum (blue dots) of network population firing rate as a function of oscillation frequency *ω*, for *α* = 1.2 (left) and *α* = 2 (right). Power-law line (orange) with exponent −2.0 provided for comparison. LFP power spectrum with power-law line of exponent −1.0 in inset.

## VI. DISCUSSION

In this study, we have developed a novel fractional diffusion theory of biologically plausible, balanced heterogeneous neural networks. By capturing the highly fluctuating, heterogeneous synaptic inputs to neurons as observed in the cortex, our theory identifies a new type of state that exhibits rich nonlinear response properties, providing a unified network mechanism underlying a great variety of neural dynamics ranging from the individual-neuron to the network levels [5, 7, 25–27], which otherwise remain disjointed in existing theories and models. Importantly, in this state the neural response is maximised as a function of structural connectivity; i.e. the firing rate for a given external input is at its greatest.

The fractional diffusion theory provides a novel theoretical framework for understanding how complex neural dynamics can emerge from heterogeneous networks. In this theory, fluctuating synaptic inputs to neurons are formulated as Lévy stochastic processes. These Lévy processes are characterized by intermittent large jumps; the presence of these processes is a hallmark of non-equilibrium complex systems [32–37]. In our theory, we rigorously demonstrate that these processes arise from heterogeneous neural networks in which the synaptic coupling weights are heavy-tailed and comparable with the size of the network. In the large network size limit, as a result of the generalized central limit theorem, this corresponds to heavy-tailed, power-law weights. This is, however, fundamentally different from recent studies of heterogeneous neural networks, in which heterogeneous connectivity has been interpreted in terms of heavy-tailed degree distributions with [18] and without [15] heavy-tailed synaptic strength distributions.

In our theory, the Lévy statistics allows for a novel application of fractional diffusion formalisms in neural networks. Previous studies of fractional diffusion in various branches of statistical physics [35] have encountered difficulties when attempting to use the fractional Fokker-Planck equation in the presence of absorbing boundaries. The integral truncations performed in those studies on the fractional Fokker-Planck operator to incorporate these boundary conditions break its convenient Fourier multiplier representation, rendering it unusable for analytical predictions. To develop an analytically tractable fractional Fokker-Planck equation with the required absorbing boundary conditions in neural networks, we construct a new link between the first passage time and the first passage leapover, previously regarded as related but independent concepts [45, 46]. Our novel fractional formalisms can thus be used to analytically reveal the circuit mechanism of complex neural dynamics, extending the recent interest in using fractional approaches for understanding complex physical systems [31, 59] to a new area.

Occasional, large synaptic inputs which are the hallmark features of Lévy processes have been found in neural circuit models, due to a combination of heavy-tailed but non-power-law connectivity features along with a specific dynamical state of the circuit [18], e.g. near a transition regime between different circuit states. We thus expect that the fractional diffusion theory applies whenever the inputs have Lévy fluctuations, regardless of specific network structures. In addition, it should be noted that scale-free, Lévy-like fluctuations can emerge from neural networks with criticality [3, 18, 60, 61]. In the future it would be interesting to apply our fractional framework for the formulation of critical neural dynamics, going beyond the demonstration of the presence of power-law distributions in some neural observables.

As we have demonstrated, our fractional theoretical framework provides a unified account of key properties of neural activity at the individual-neuron and the circuit levels. First, membrane potentials in our model exhibit non-Gaussian dynamics, consisting of quiescent periods interrupted by short intervals of high-amplitude depolarization, and reside far from the threshold. Such non-Gaussian fluctuations of membrane potential have been observed in the spontaneous activity of cortical neurons [27] but are inconsistent with the existing random walk models [62], which instead predict Gaussian dynamics of membrane potential. Large fluctuations of membrane potential could be obtained from classical excitatory-inhibitory balanced networks with a suitable choice of parameters, leading to Gaussian statistics with frequent fluctuations on the order of the standard deviation. However, the infrequent, occasional large spontaneous fluctuations observed in [27] cannot be compatible with Gaussian input statistics, as noted in [27], but can be captured in our model. Moreover, while the classical excitatory-inhibitory spiking circuits have been able to account for the fluctuations of up- and down-states of membrane potentials, which have often been observed in cortex of anesthetized animals, by incorporating additional mechanisms such as adaptation [28], the fractional diffusion theory is mainly used in this study to explain the brief, large excursions of membrane potential dynamics observed in awake cortex [27, 63]. The non-Gaussian state in [27], which occurs during awake fixation, is characterized by infrequent, brief bumps in inputs, which constitute a defining feature of the Lévy process.

Second, the variance of spike counts is a super-linear function of their mean as measured in [7], and the Fano factor of spike counts increases as a power-law as a function of the time window, as found in [24, 26]. In contrast, in the classical normal diffusion theory, the variance of spike counts is equal to the mean, and the Fano factor remains constant over large timescales. Phenomenological models based on the Poisson process with the incorporation of global fluctuating modulation factors have been proposed to account for such fluctuations of firing activity [7]. Because the sources of these global fluctuations are unknown, these phenomenological models are somewhat circular, leaving the fundamental problem of the circuit mechanism underlying the fluctuations unaddressed. By demonstrating that such firing rate fluctuations are an intrinsic property of our heterogeneous balanced circuits, our study thus identifies a novel circuit mechanism by which these phenomena can be explained.

Third, firing rates across neurons are highly heterogeneous, with some neurons firing a lot but others firing much less, resulting in the heavy-tailed lognormal distribution as widely observed in experimental studies [23]. Fourth, the population firing rate and local field potentials in the fractional state oscillate across multiple frequency scales in a 1*/f* manner, and all oscillations, such as gamma oscillations at 40 Hz, occur intermittently in a bursting manner, consistent with experimental studies of population activity recorded by local field potentials [25, 39]. Achieving such nonlinear behaviors of oscillations in conventional balanced neural networks has remained a major challenge, due to the presence of a single characteristic frequency as predicted by the linear stability analysis in the conventional Fokker-Planck formalism [10, 38].

Our theory identifies an optimal state of activity, in which the response of the circuit to input is nonlinear and structurally optimized. Due solely to the heterogeneity of the network, the neural response for the same average input is significantly higher than that of the asynchronous state as in the classical framework based on the Gaussian assumption, and in the optimal state, the stationary firing rate is maximal as a function of the fractional Lévy index *α* arising in the synaptic strength distribution. In the fractional regime, all physiological parameters of the network, such as the excitatory-inhibitory balance, the external input size, the mean connection strength, or the mean in-degree, have remained unchanged from the classical case. Furthermore, in the fractional regime, other variability properties are consistent with physiological findings; this drives us to use the term ‘regime’ to capture a range of nonlinear, non-equilibrium phenomena which cannot be obtained by changing other constraints. In this context, a stronger response for balanced networks based purely on the connectivity structure of the network, without changing physiological, magnitude-based properties, becomes of fundamental computational importance, and can thus link structure with dynamics. This optimal response uncovered in our fractional theory may have important functional consequences for the biological requirements of neural efficiency. To test this structurally optimal regime with the essentially nonlinear computation properties revealed in our study, it would be ideal to combine intra- and extra-cellular recordings with massive multi-unit recordings in order to measure the fluctuations of membrane potentials and firing activity at the individual neuron and population levels. This could be done in conjunction with the analysis of membrane potential, synaptic input and neural fluctuations by using the same methods as we have done in our modeling study.

### Appendix A: Derivation of Lévy stable input

To demonstrate the emergence of Lévy stable input, we consider a single neuron in a network undergoing a discrete time evolution with timestep Δ*t*, so the probability of a neuron in population *r* being active at a given timestep is ⟨*s*(*t*) ⟩ _*r*_ := *v*_*r*_(*t* − *D*)Δ*t*. The probability that a neuron receives a connection from a given neuron in population *r* is *C*_*r*_*/N*_*r*_. The general idea here is to consider the inputs from each population of incoming neurons separately, and use the properties of stable processes to combine them into the total input. The outgoing connection strengths from population *r* are drawn from a distribution characterizing the random variable *J*_*r*_. If *X*_*r*_ is the random variable representing the firing of an incoming neuron from population *r*, which is Bernoulli with probability *λ*_*r*_*/N*_*r*_, where *λ*_*r*_ := *C*_*r*_ ⟨*s*(*t*) ⟩ _*r*_, then the characteristic function of *J*_*r*_*X*_*r*_ is

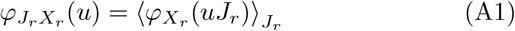

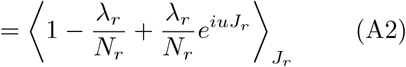

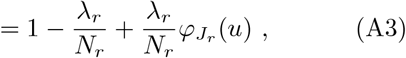

generalizing the characteristic function of a Bernoulli random variable. Suppose now that *X*_*i*_, *J*_*i*_ are independent and distributed as *X*_*r*_, *J*_*r*_, for all *i* = 1, *…, N*_*r*_. The characteristic function for the total synaptic input to the neuron from population *r* is

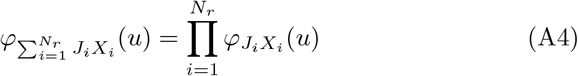

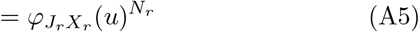

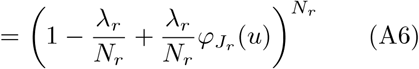

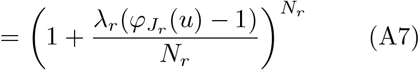

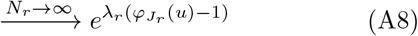

generalizing the Poisson limit theorem. Then, using Lévy’s continuity theorem [43],

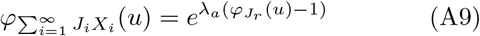

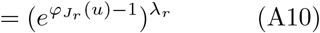

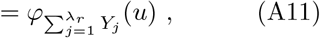

where *λ*_*r*_ is taken to be an integer in the last equality (valid for large *λ*_*r*_), and the *Y*_*j*_ are iid random variables with 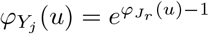. The cumulant generating function (second characteristic function) of *Y*_*j*_ is

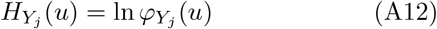

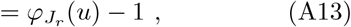

so the cumulants of *Y*_*j*_ are precisely the moments of *J*_*r*_; in particular, ⟨*Y*_*j*_ ⟩= ⟨*J*_*r*_⟩.

The remainder of the derivation is split into two cases: when the second moment (or equivalently, the variance) of *J*_*r*_ is infinite, and when it is finite. We consider first the former case. Suppose that the random variable *J*_*r*_ satisfies

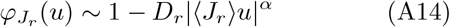

as *u* → 0, so that its probability density function has a positive tail of the form

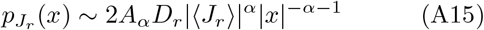

as *x* + → ∞, where *A*_*α*_ = Γ(1 + *α*) sin(*πα/*2)*/π* [64, p10-11] is a constant independent of the population, and a negative tail which decays more rapidly. Then, using the generalized central limit theorem [43],

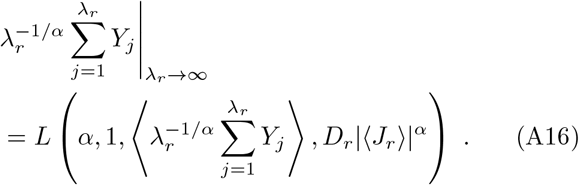

If, instead, the connection strengths of the population have a dominant negative tail (e.g. the inhibitory population), the above holds with the exception of the skew parameter *β*_*r*_ taking the value −1. Hence, using the additive property of the stable law, the total synaptic input to the neuron from all populations is

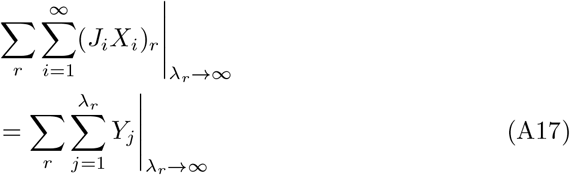

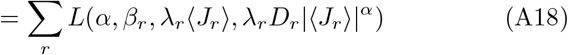

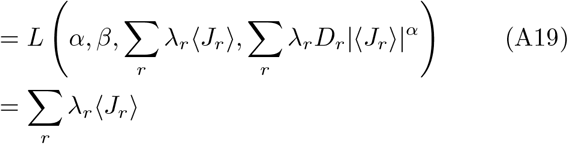

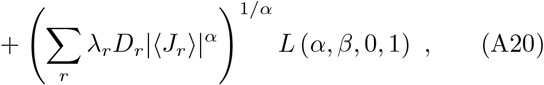

where 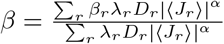. Moving into the continuous-time limit, one obtains

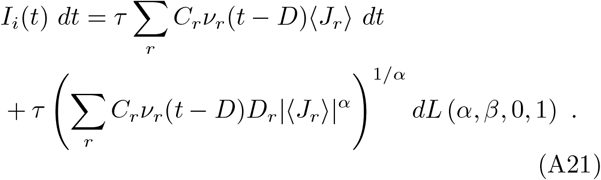

If all populations have the same coefficient *D*_*r*_ = *D*_*L*_, then

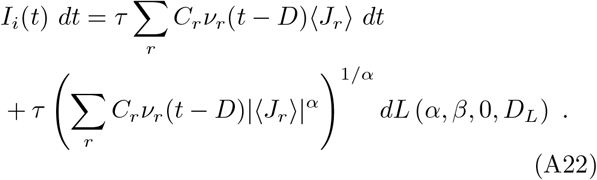

Since *C*_*I*_ = *γC*_*E*_, the skew parameter of the noise in our excitatory-inhibitory network model becomes

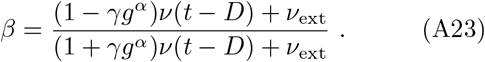

In the latter case, when *J*_*r*_ has finite second moment, the generalized central limit theorem reduces to the regular central limit theorem and *α* = 2. Choose *D*_*r*_ such that

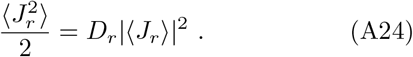

Since the second moment of *J*_*r*_ is equal to the second cumulant (the variance) of *Y*_*j*_, the above derivation for 1 *< α <* 2 extends to the *α* = 2 case with Eqs. (A14, A15) replaced by Eq. (A24).

The generalized central limit theorem, in the form stated in Table 2.1 of [43], allows for another case (not investigated in this study). When 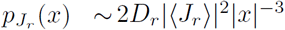 (note the absence of the *A*_*α*_ term) the resultant dynamics of the model become Gaussian again, with the sole change being the replacement of the *x*_*r*_ := *C*_*r*_*v*_*r*_(*t* − *D*_*B*_) term in *σ*(*t*) with *x*_*r*_ log *x*_*r*_.

**TABLE 1.** Parameter sets used in the main text.

| | $\alpha = 1.2$ | $\alpha = 2$ | Units |
| --- | --- | --- | --- |
| $N_E$ | 40 000 | 10 000 | |
| $N_I$ | 10 000 | 2 500 | |
| $\gamma$ | 0.25 | 0.25 | |
| $C_E$ | 4000 | 1000 | |
| $C_I$ | 1000 | 250 | |
| $C_{\text{ext}}$ | 4000 | 1000 | |
| $\tau$ | 20 | 20 | ms |
| $D$ | 1.5 | 1.5 | ms |
| $\nu_{\text{ext}}$ | 1 | 2 | $\nu_{\text{thr}}$ |
| $\theta$ | 20 | 20 | mV |
| $V_r$ | 10 | 10 | mV |
| $\tau_{\text{rp}}$ | 2 | 2 | ms |
| $J$ | 0.02 | 0.1 | mV |
| $g$ | 4 | 5 | |

### Appendix B: Lévy stable input with heterogeneous firing rate distributions

In this section we demonstrate how the fractional mean-field theory in Appendix A may be extended to predict distributions of population firing rates. In the classical theory, where the firing rate, synaptic strength, and number of connections all had finite variance across the network, this involved adding a quenched Gaussian term to the mean input 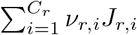 for each population *r*. This corresponds to the first term on the right-hand sides of Eqs. (A20,A21). Due to the Gaussian randomness, it was sufficient to perform this to first order for each of these sources of randomness [12–14, 65].

In the fractional diffusion theory, the same technique would yield a firing rate distribution describing the network in the absence of the tight balance condition (23). However, incorporating this balance condition causes the first-order terms to cancel out for the internal populations, and the analysis for these populations must then be brought to second order.

Using the generalized central limit theorem and the synaptic heavy-tailed assumption (A15), the mean input arriving from population *r* to a randomly selected network neuron approaches, in the limit of a large number of connections,

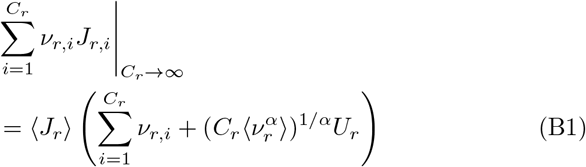

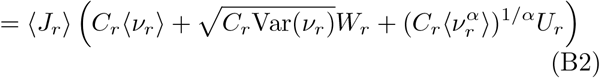

where *v*_*r,i*_ are samples drawn from the population firing rate distribution *v*_*r*_; *U*_*r*_ ∼ *L*(*α*, 1, 0, *D*_*r*_) is a quenched *α*-stable sample; and *W*_*r*_ ∼ 𝒩(0, 1) = *L*(2, 0, 0, 1*/*2) is a quenched Gaussian sample. When *C*_*r*_ is large, the central limit theorem applies on 0 ≤ *v*_*r*_ ≤ 1*/τ*_rp_ since it is bounded. This yields a closed, self-consistent system of equations to second order in the absence of tight balance.

To incorporate the balance condition (23), we observe that the above implies for the inhibitory population that

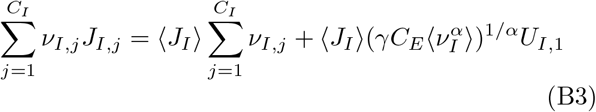

in the limit of large *C*_*I*_. Similarly, taking another *C*_*E*_ new samples of the product *v*_*I*_ *J*_*I*_,

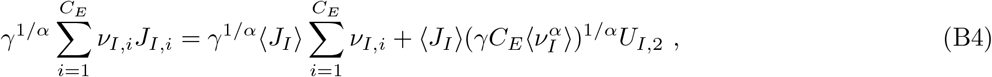

and we find that in distribution

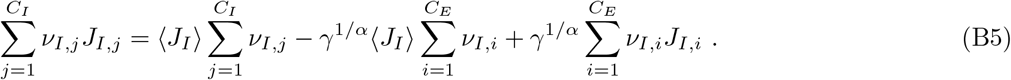

The tight balance condition (23) then allows us to take *J*_*I,i*_ = −*gJ*_*E,i*_, noting that *J*_*I,i*_ are random samples of the distribution *J*_*I*_ which do not appear as network synaptic strengths; similarly the *v*_*I,i*_ do not appear in the network as individual neural firing rates. As a result, we have that in distribution the mean input from the internal populations with tight balance is

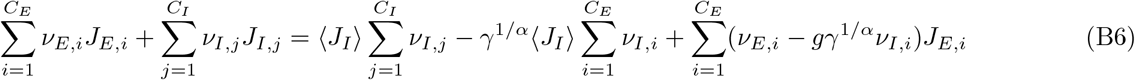

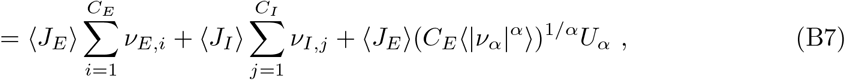

where

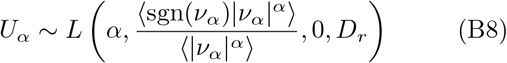

is a quenched *α*-stable sample, and 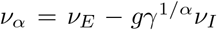 is a random variable. Since in our model excitatory and inhibitory neurons have identical characteristics, the distributional equality *v*_*E*_ = *v*_*I*_ yields the following modification to Eq. (3) for the input *I*(*t*) to a randomly selected neuron at time *t*:

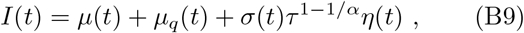

where

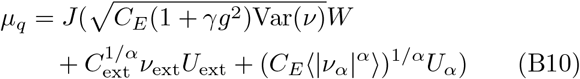

is the quenched contribution to the mean input, and instances of *v*(*t*) are replaced with their population means, ⟨*v*⟩ (*t*). This system of equations is closed through the definition of ⟨*v*⟩ (*t*), Var(*v*)(*t*), ⟨|*v*_*α*_|^*α*^⟩ (*t*) and ⟨sgn(*v*_*α*_)|*v*_*α*_|^*α*^⟩ (*t*).

Finally, the above is conducted under the assumption that the firing rate of a neuron is uncorrelated with its outgoing synaptic strengths. This may no longer be the case in such strongly connected networks, especially given the propensity for networks with rich clubs to exhibit local rather than global behavior [18]. We have attempted to account for this by assuming that neurons with low firing rates have a small sum of incoming synaptic strength magnitudes, and vice versa. However, the mean-field construction links the firing rate of a neuron with only its incoming, rather than both its incoming and outgoing synaptic strengths. This approaches the limit of what can be achieved using such mean-field techniques: the correlations between the outgoing and incoming properties of individual elements begin to become too significant for a general mean-field analysis. Unlike the situation in [13], where the correlations between neural firing rates and their *a priori* correlated in- and out-degrees were able to be considered, the incoming and outgoing synaptic strengths in our model are independent. In the future, new techniques, possibly inspired by fractal, renormalization group-like methods, may replace the mean-field technique in describing the behaviour of heterogeneous, complex systems in statistical physics.

### Appendix C: Derivation of the stationary states

Here we present the technical derivation of the stationary states of the system, which consists of solving a fractional differential equation. Classical analyses of FFPEs in the absence of absorbing boundary conditions tend to prefer solving the system in Fourier-Laplace space [66], where the relevant dynamical operators have a simple form. Using our analytically tractable absorbing boundary conditions, we undertake a similar method whereby we move into Fourier space, and take the inverse Fourier transform of the solution to the Fourier ordinary differential equation.

In the stationary state (where *q*_0_(*V*) := *q*(*V*, 1*/v*_0_)),

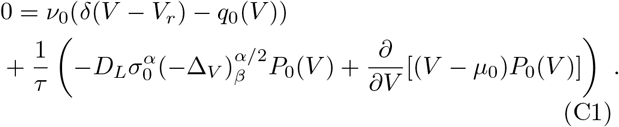

With the aid of the following rudimentary properties of the Fourier transform of a function *f* ∈ ℒ^*p*^(ℝ),

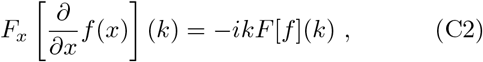

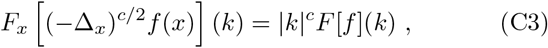

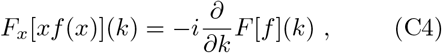

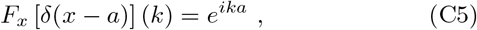

where *p* ∈ [1, 2] and *c* ∈ (0, 2], Eq. (C1) is expressed in Fourier space as

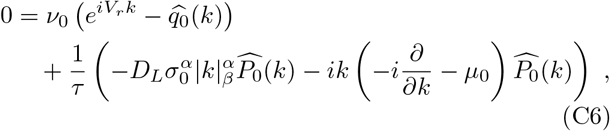

where

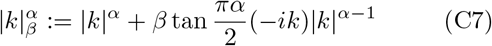

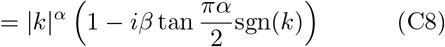

is the multiplier corresponding to the skewed fractional Laplacian 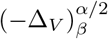. The solution to this first-order differential equation is

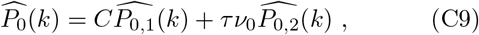

where

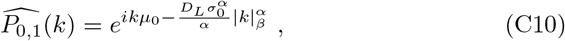

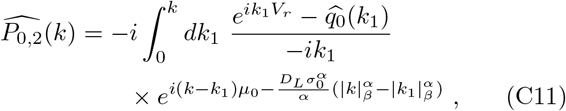

and *C* is a constant determined by the boundary conditions, and so

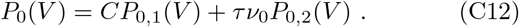

Taking the inverse Fourier transform of Eqs. (C10, C11) yields expressions in terms of the standard alpha-stable distribution,

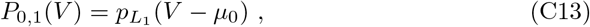

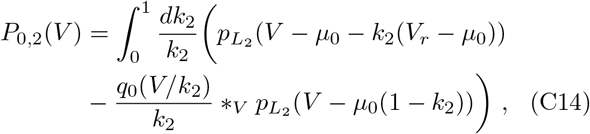

Where

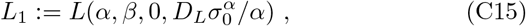

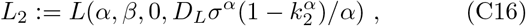

and *f* (*V*) *_*V*_ *g*(*V*) = ∫*f* (*V* − *V*_1_)*g*(*V*_1_)*dV*_1_ is the convolution operator. To understand the effect of the threshold *θ* on the solution, Eq. (C14) can be written as

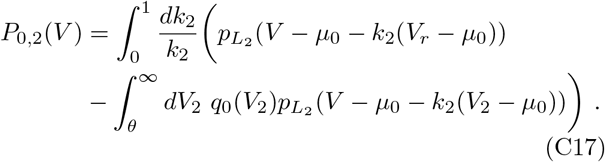

To increase computational accuracy, the boundaries in the inner integral in Eq. (C17) can be rewritten over ℝ, giving

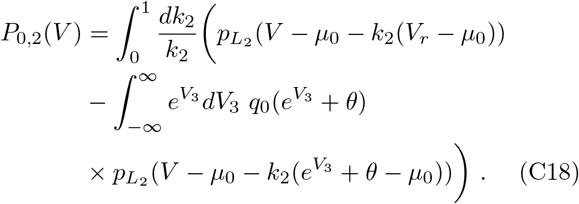

Using the two boundary conditions (17), (19) to determine the two remaining undetermined variables *v*_0_ and *C*, one obtains 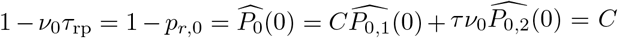 and 0 = *P*_0_(*θ*) = *CP*_0,1_(*θ*) + *τv*_0_*P*_0,2_(*θ*). Thus the self-consistent condition that determines *v*_0_, the stationary firing rate, is

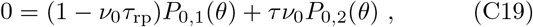

or equivalently

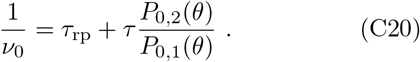

### Appendix D: Network implementation details

To visualize the behavior of the system in this study, it is necessary to specify biologically realistic numerical values for the network parameters. These values are presented for the two parameter sets used in the main text in Table I. The biologically realistic values are derived from the balanced case in Section 4 of [10], with a ten-fold decrease in the mean excitatory connection strength *J* in order to produce realistic membrane potential distributions in the Gaussian regime. The parameters for the classical, asynchronous irregular case (*α* = 2) are derived from Fig. 7(c) of [10].

To construct a computationally tractable synaptic strength distribution for the network satisfying Eq. (A15), we consider the lower-bound-truncated translated power-law distribution for the synaptic connection strengths,

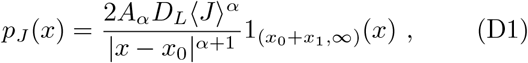

where *x*_0_, *x*_1_ is determined by the normalization condition 1 = ⟨1⟩, and the constraint on the first moment of *J* is ⟨*J*⟩ = ⟨*J* − *x*_0_⟩ + *x*_0_. Using the relation

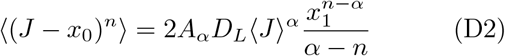

we obtain

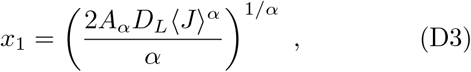

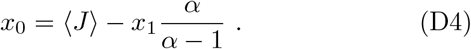

One can then sample values from this distribution using NumPy’s pareto function.

For comparison with previous analyses [10, 38], although the exact value of the common scale *D*_*L*_ depends on the shape of the synaptic strength distribution *J*_*r*_, we have chosen it in computations to coincide, when *α* = 2, with the diffusion approximation case where the synaptic strengths are constant for all neurons in each population. In this classical case, |⟨*J*⟩| ^2^ = ⟨*J*⟩^2^, and so *D*_*L*_ = 1*/*2 using Eq. (A24).

